# Comparing short and long batteries to assess deficits and their neural bases in stroke aphasia

**DOI:** 10.1101/2020.11.24.395590

**Authors:** Ajay D. Halai, Blanca De Dios Perez, James D. Stefaniak, Matthew A. Lambon Ralph

**Author notes:** Joint first authors (Ajay D. Halai, Blanca De Dios Perez). Corresponding authors Dr. Ajay Halai or Prof. Matthew Lambon Ralph MRC Cognition & Brain Sciences Unit, University of Cambridge, 15 Chaucer Road, Cambridge, CB2 7EF. United Kingdom.

## Abstract

Multiple language assessments are necessary for diagnosing, characterising and quantifying the multifaceted deficits observed in many patients’ post-stroke. Current language batteries, however, tend to be an imperfect trade-off between time and sensitivity of assessment. There have hitherto been two main types of battery. Extensive batteries provide thorough information but are impractically long for application in clinical settings or large-scale research studies. Clinically-targeted batteries tend to provide superficial information about a large number of language skills in a relatively short period of time by reducing the depth of each test but, consequently, can struggle to identify mild deficits, qualify the level of each impairment or reveal the underlying component structure. In the current study, we compared these batteries across a large group of individuals with chronic stroke aphasia to determine their utility. In addition, we developed a data-driven reduced version of an extensive battery that maintained sensitivity to mild impairment, ability to grade deficits and the component structure. The underlying structure of these three language batteries (extensive, shallow and data-reduced) was analysed using cross-validation analysis and principal component analysis. This revealed a four-factor solution for the extensive and data-reduced batteries, identifying phonology, semantic skills, fluency and executive function in contrast to a two-factor solution using the shallow battery (phonological/language severity and cognitive severity). Lesion symptom mapping using participants’ factor scores identified convergent neural structures based on existing language models for phonology (superior temporal gyrus), semantics (inferior temporal gyrus), speech fluency (precentral gyrus) and executive function (lateral occipitotemporal cortex) based on the extensive and data-reduced batteries. The two components in the shallow battery converged with the phonology and executive function clusters. In addition, we show that multivariate prediction models could be utilised to predict the component scores using neural data, however not for every component score within every test battery. Overall, the data-reduced battery appears to be an effective way to save assessment time yet retain the underlying structure of language and cognitive deficits observed in post stroke aphasia.

## 1. Introduction

It is critical to have accurate and reliable ways of measuring symptoms, in order to perform differential diagnosis and implement the optimum treatment pathway. For neuropsychological disorders, the issue of measuring symptoms is non-trivial for a number of reasons. First, patients can have a wide range of deficits (e.g., memory, attention, speech and language, etc.), thus potentially necessitating a large number of assessments. Second, any given test needs sufficient dynamic range to capture a wide range of severities (complete impairment to well-recovered), which requires a sufficient number of items with varying degrees of difficulty to avoid floor or ceiling effects. This is particularly important when deficits are graded along principal behavioural axes as opposed to falling into classic binary distinctions (Lambon Ralph *et al.*, 2003; Butler *et al.*, 2014). Capturing the full range of deficits and their entire severity range requires an extensive, detailed assessment battery, which is rarely feasible in clinical settings, large-scale clinical trials or where patients have attention/fatigue deficits. The current study explored this challenging issue and the efficacy of alternative assessment strategies through the test case of post-stroke aphasia. Diagnosing language and cognitive deficits in post-stroke aphasia is particularly challenging as there is considerable variation in the cognitive/language domains affected and the severity of the impairments. In order to save time, most batteries adopt a “shallow” approach, i.e., preserve the breadth (test many domains) but reduce the depth of each test (number of items). In the current study we directly compared an extensive battery (containing numerous tests each with many assessment items) against (a) a popular ‘shallow’ battery, the Comprehensive Aphasia Test (CAT) (Swinburn *et al.*, 2004); and (b) a novel data-driven ‘reduced’ test battery which limited the number of tests included but preserved their “depth”. For each, we investigated their ability: (i) to detect and grade the patients’ impairments; (ii) to reveal the underlying principal dimensions of variations across the patient cohort; and (iii) to map the corresponding lesion correlates.

The long history of aphasia research contains many different approaches to assessment including early examples of systematic test batteries (Head, 1920). Many famous, popular batteries were designed to provide efficient clinical diagnoses of aphasia and their subtypes (i.e. Boston Diagnostic Aphasia Examination [BDAE] (Goodglass *et al.*, 1972), Western Aphasia Battery [WAB] (Kertesz, 1982), Minnesota test for differential diagnosis of aphasia [MTDDA] (Schuell and Sefer, 1965), Porch Index of Communicative Ability [PICA] (Porch, 1967)). Many of these, however, have been found to be inadequate at identifying the nature of language impairments and guiding future interventions (Byng *et al.*, 1990). Alternative approaches included batteries in the form of a ‘bank’ of psycholinguistically-sophisticated and detailed tests, such as the Psycholinguistic Assessment of Language Processing in Aphasia (Kay *et al.*, 1992), from which experts select assessments to suit each individual patient. More recently, this style of psycholinguistically-informed tests were transformed into a new ‘shallow’, systematic battery (the Comprehensive Aphasia Test: (Howard *et al.*, 2010)). The CAT is usually administered over 1-2 hours and contains three sections: 1) cognitive screening; 2) language battery; and 3) a disability questionnaire. The language battery probes many different language activities each with a minimum number of carefully chosen items. The CAT was always intended to be an initial screening battery to be followed up by more detailed assessment of the identified areas of interest for each patient. Unsurprisingly, this efficient battery is used both clinically and in numerous research projects.

A second core aim of the current study was to examine the ability of different types of assessment battery to capture the underlying variations in post-stroke aphasia (PSA). The considerable inter-participant variations in PSA are well known as are the limitations of considering these differences in terms of categorical classifications, which fail to capture important aspects about the underlying impairments, and are unable to relate classifications and the underlying lesions (Poeck, 1983; Basso, 2003; Howard *et al.*, 2010). Based on detailed assessment batteries, contemporary studies have begun to reconceptualise PSA in terms of graded variations along a limited number of underpinning principal language and cognitive dimensions (e.g., phonology, semantics, fluency and executive-cognitive skill), each of which is clearly associated with specific critical brain regions (Butler et al., 2014; Halai et al., 2017; Lacey et al., 2017; Mirman, et al., 2015a; Mirman, et al., 2015b). Interestingly, similar analyses have been conducted on each section of the CAT separately (Swinburn *et al.*, 2004). One dimension was obtained after applying PCA to the cognitive screen subtests, onto which all tests loaded strongly except line bisection. The language tests collapsed into three factors: comprehension (and writing), repetition and reading. The first two components could reflect the semantics and phonology factors found in the recent large-scale examinations noted above. Reading from the CAT might also span these same two components, as a recent large-scale study has implicated nonword reading with phonological abilities, whilst word reading calls upon phonology and semantics in tandem (Woollams *et al.*, 2018). Key questions, therefore, for the current study included: (a) how well can different types of battery (full, shallow, reduced – see next) reveal the full collection of underlying dimensions; and (b) what dimensions are revealed by the CAT battery when the language and cognitive measures are analysed simultaneously.

The use of PCA and other data-reduction techniques are also relevant to the current study for another reason. One of the first studies of PCA in PSA (Butler *et al.*, 2014), found that it was possible to use the PCA task loadings to identify which individual tests best approximate each underlying dimension. We used this finding as the basis for generating a different kind of reduced battery. Specifically, principal component analysis was used to determine: 1) which subset of tests are the best proxies for each principal component; and 2) within each test, which subset of items best capture the variance in that test’s data. By applying this method to the extensive battery, we generated a data-driven ‘reduced’ battery that is quick and efficient to administer, yet retains the extensive battery’s sensitivity for the underlying component structure.

Finally, we examined the ability of each type of battery identify the corresponding neural correlates. In previous work, we mapped the four principal components to the integrity of discrete brain regions (Halai *et al.*, 2017) that align with results from fMRI language studies in healthy participants (e.g. Hickok & Poeppel, 2007; Price, 2012). A number of studies have mapped different subsets of the CAT to brain damage (e.g., Hope *et al.*, 2013, 2015, 2018). To gain a complete picture, in the current study we compared neural correlates that arise from each of the three batteries. Lesion-symptom mapping can now be conducted using univariate or multivariate methods (Bates *et al.*, 2003; Tyler *et al.*, 2005; Mah *et al.*, 2014; Zhang *et al.*, 2014; DeMarco and Turkeltaub, 2018; Sperber and Karnath, 2018). Although there are strong advocates for each one, these alterative analyses tackle different fundamental questions, and have opposite strengths and weaknesses (Schumacher *et al.*, 2019). Multivariate methods are predictive in nature and account for co-dependencies between features. This means, though, that obtaining local inference is inherently difficult as the models rely on a combination of (usually distributed) beta weights, which cannot be thresholded post-hoc (i.e. using permutation testing) unless a feature selection strategy or sparse solution is implemented. Furthermore, the beta weights assigned to features are not transparent (Haufe *et al.*, 2014; Hebart and Baker, 2018) and therefore caution must be exercised before making strong inferences about high/low weights. The opposing strengths and weaknesses are true for the univariate approaches, where local inferences and interpretation of weight strengths are straightforward yet such approaches might miss key dependences between regions and/or mislocalise the true effect (Mah *et al.*, 2014; Zhang *et al.*, 2014; Sperber and Karnath, 2018). With these issues in mind, in the current study we present both univariate and multivariate analyses for each test battery.

## 2. Materials and Methods

### 2.1. Participants

Seventy-five chronic post-stroke (haemorrhagic or ischaemic) patients with aphasia were recruited for this study. Participants were assessed with the short form of the BDAE and assigned an aphasia classification (Goodglass *et al.*, 1972). All participants were at least twelve months post-stroke, native English speakers with normal or corrected-to-normal hearing and vision. Participants were excluded based on the following criteria; having more than one stroke, other neurological conditions, contraindications for MR scanning or being left handed premorbidly. All cases had extensive neuropsychology and neuroimaging assessments (detailed below); additionally, a subgroup (N = 40) completed the CAT.

The demographic characteristics are presented in Supplementary Materials Table 1. Informed consent was obtained from all participants prior to participation in the study under approval from the local ethics committee.

**Table 1.**
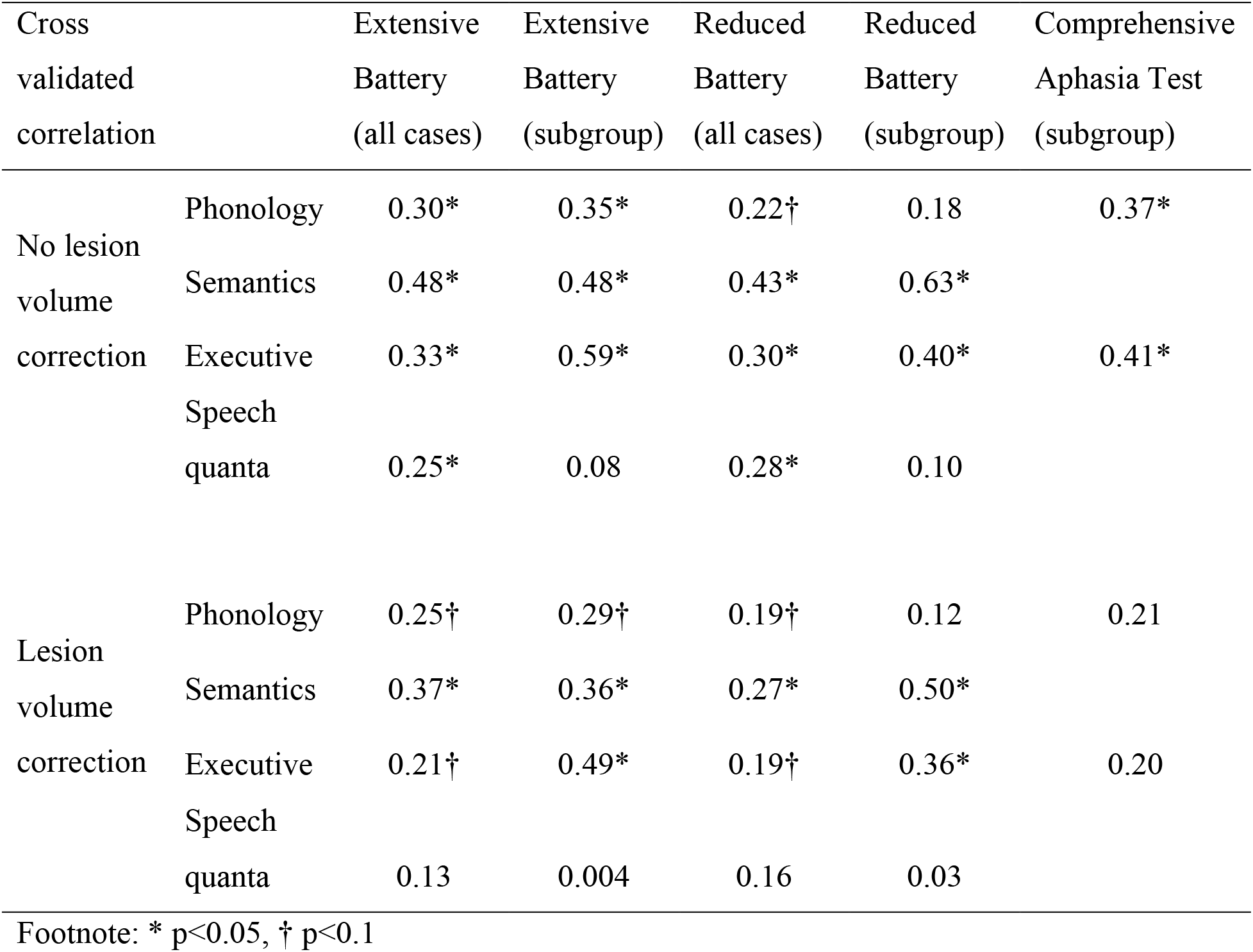
Results from multivariate models predicting each principal component from brain abnormality images. The table shows the cross validated correlation between predicted and observed scores, where significant models were determined using permutation testing (N = 1,000). Model performances with and without lesion volume correction are shown.

### 2.2. Assessment

All participants were tested on an extensive neuropsychological battery described in (Butler *et al.*, 2014; Halai *et al.*, 2017). The battery included several subtests from the Psycholinguistic assessment of language processing in aphasia (PALPA) (Kay *et al.*, 1992): immediate and delayed repetition of non-words (PALPA 8); immediate and delayed repetition of words (PALPA 9). Tests from the Cambridge Semantic Battery (Bozeat *et al.*, 2000) included: spoken and written word-to-picture matching; 64-item picture naming task; and Camel and Cactus with pictures (CCTp). We also included the Boston Naming Test (BNT) (Goodglass *et al.*, 1972), the 96-item synonym judgement test (Jefferies *et al.*, 2009), comprehension of spoken sentences from the CAT, and forward and backward digit span (Wechsler, 1987). We also included cognitively demanding nonverbal tests, the Brixton Spatial Rule Anticipation Task (Burgess and Shallice, 1997) and Raven’s Coloured Progressive Matrices (Raven, 1962). Four measures of fluency were extracted from the BDAE ‘Cookie Theft’ picture description task (Goodglass *et al.*, 1972): number of speech tokens (T), words per minute (WPM), mean length per utterance (MLU) and type token ratio (TTR) (details of coding are provided in Borovsky, Saygin, Bates, & Dronkers, 2007 and Halai et al., 2017).

All participants completed the extensive battery first as part of a larger on-going data collection protocol. We successfully re-visited forty participants to assess their performance on the CAT (electronic version) (Swinburn *et al.*, 2004) omitting the disability questionnaire section as it was not relevant to the current study.

### 2.3. Reduced Battery

Our goal was to reduce the time it would take to administer neuropsychological testing while retaining sensitivity to the underlying component structure. To determine this target structure we took the extensive neuropsychological test battery in our full cohort of 75 patients and applied a varimax rotated principal component analysis (SPSS v20.0). For each principal component with an eigenvalue greater than 1 (the optimal number of components were also confirmed using k-fold cross-validation, see below for details), we included two tests in the reduced battery as representative proxies. Specifically, we took tests that loaded high on the target dimension and near zero on others as well as constraining selection with our knowledge of their clinical utility (i.e., if there were multiple high loading tests, we took the test that would be easiest to administer in a clinical setting). We excluded tests if they loaded onto multiple principal components (with a loading score >0.5). The only exceptions were the tests of naming and sentence comprehension because these are functionally important tasks for patients to be able to perform irrespective of their relationship to the underlying component structure of language. We therefore included the BNT, Cambridge Semantic Battery 64-item picture naming test and CAT spoken sentence comprehension test.

As well as reducing the number of tests, we also sought to reduce the number of items in some of the longer assessments. For instance, the PALPA9, BNT, Cambridge Semantic Battery, and synonym judgement tests contained over 60 items each. We therefore halved the number of items in these tests in a data-driven manner. To achieve this, we coded item level responses for each of the 75 PSA participants for each test and performed an unrotated factor analysis restricted to a one factor solution. The top 50% of items loading most strongly on the identified factor were included in the reduced item set for each test. Certain tests had an internal structure (i.e. factorial design) that respected psycholinguistic distinctions: the 96-item synonym judgement test manipulates word frequency (2 levels) and imageability (3 levels) yielding 6 distinct classes, while the Cambridge Semantic Battery 64-item picture naming comprised 32 living and 32 non-living items. For these tests, we conducted a separate factor analysis on each factorial level to retain the internal structure. Further details of the reduced tests are shown in Supplementary Materials Section 2.

### 2.4. K-Fold Cross Validation Analysis

In order to check the stability and reliability of the PCA solutions, we performed five-fold cross-validation analyses (Ballabio, 2015) (version 1.3 in MATLAB 2018a). This procedure allows us to determine the optimum number of components in our dataset by performing a PCA on a training set and predicting the scores of left-out cases (based on venetian blinds sampling). The prediction is carried out for N-1 models to determine which number of components provides the best solution corresponding to the lowest root mean squared error (N = number of tests). The behavioural data were scaled to percentage and the training data were normalised to z-scores before submitting to the cross-validation analysis. Once an optimal number of components was determined we performed a second leave-one-out validation analysis. In this analysis, a model was created using the optimal component number on the training data and the test data were predicted (by projecting the left-out data into the trained component space using the coefficient matrix). A correlation was obtained between the observed and predicted data as a measure of generalisability of the PCA model.

### 2.5. Acquisition of neuroimaging data

High resolution structural T1-weighted Magnetic Resonance Imaging (MRI) scans were acquired on a 3.0 Tesla Philips Achieva scanner (Philips Healthcare, Best, The Netherlands) using an eight-element SENSE head coil. A T1-weighted inversion recovery sequence with 3D acquisition was employed, with the following parameters: TR (repetition time) = 9.0ms, TE (echo time) = 3.93ms, flip angle = 8°, 150 contiguous slices, slice thickness = 1 mm, acquired voxel size 1 × 1 × 1 mm^3^, matrix size 256 × 256, FOV= 256 mm, TI (inversion time) = 1150ms, and SENSE acceleration factor 2.5 with a total scan acquisition time of 575 s.

### 2.6. Analysis of neuroimaging data

Structural MRI scans were pre-processed with Statistical Parametric Mapping software (SPM12: Wellcome Trust Centre for Neuroimaging, https://www.fil.ion.ucl.ac.uk/spm/). Images were normalised into standard Montreal Neurological Institute (MNI) space using a modified unified segmentation-normalisation procedure optimised for focally lesioned brains (Automated Lesion Identification – ALI v3) (Seghier *et al.*, 2008). The resulting lesion outputs were visually inspected for accuracy and manually adjusted if needed.

We conducted univariate and multivariate brain-behaviour mapping using the PCA component scores derived from: 1) the extensive test battery; 2) the data-driven reduced test battery; and 3) the CAT. Both brain-behaviour mapping approaches utilised the abnormality images from the ALI toolbox (hypo-intensity changes only, where each voxel is compared to a group of age and education matched controls and assigned a probability of abnormality). In the univariate analyses, we created three models (one for each PCA solution) and entered the corresponding components simultaneously. Voxel based correlational methodology (VBCM) (Tyler *et al.*, 2005) was implemented in SPM12 to determine significant clusters, using a voxelwise threshold p<0.001 with a family-wise error corrected (FWEc) clusterwise threshold p<0.05. For transparency we calculated the model with and without lesion volume as an additional covariate. Lesion volume was obtained through the automated lesion identification method (Seghier *et al.*, 2008). Anatomical labels used in the report are obtained from the Harvard-Oxford cortical and subcortical atlas and John Hopkins University white matter atlas in MNI space. We used the pattern recognition of neuroimaging toolbox (PRoNTo V2.1; http://www.mlnl.cs.ucl.ac.uk/pronto/) (Schrouff *et al.*, 2013) to determine whether individual scores on principal components could be predicted based on multivariate analysis of the abnormality detected in the T1 image. We performed the regression analysis using the relevance vector regression (Tipping, 2001) on a masked region defined by thresholding the lesion overlap map at 10%. We chose this method as it is computationally efficient compared to other machines available in the package, which makes permutation testing of a large number of models more feasible. PRoNTo uses kernel methods to minimize the high dimensionality problem, where a pair-wise similarity matrix is built between all neuroimaging scans (mean centred). The implementation does not require hyperparameter optimisation and all models were assessed for performance using a leave-one-out cross-validation scheme (k-fold was not used due to small sample size). Model inference was determined by permutation testing (N = 1,000), where the dependant variable was shuffled randomly and the permuted correlations were used as the null distribution (alpha p < 0.05).

## Data availability

Data are potentially available by request to M.A.L.R

## 3. Results

### 3.1 Patient demographics and lesion overlap

There were no significant differences (p’s.> 0.05) between the full and subgroup participants in: age (62.59 [SD = 11.43] and 62.95 [SD = 11.56] years, respectively), education (12.04 [SD = 2.10] and 12.33 [SD = 2.37] years, respectively) and months post stroke (55.51 [SD = 48.22] and 52.08 [SD = 50.32] months, respectively). The gender composition of the groups was also not significantly different (55/20 and 27/13 males and females, respectively).

We compared the lesion and behavioural profile of patients between the full and sub group. The top panels in Figure 1 show the lesion distribution for all participants and the subgroup. This primarily covers the areas of the left hemisphere supplied by the middle cerebral artery. We performed a Fischer exact test at each voxel across the brain to determine if the proportion of intact/damaged cases differed between the groups and found no significant differences (voxelwise p’s > 0.12), suggesting that the lesion profile was similar between groups. Furthermore, the lesion volume was not different between the full and sub group (16809 [SD=11555] and 16230 [SD=11493] number of voxels, respectively). In terms of behavioural profiles, rather than compare all raw test scores we compared the principal component scores (described in Section 3.3) for the full and sub groups extracted from the largest dataset available. Again, we found no significant differences between groups for any component (p’s > 0.27). The lower panel of Figure 1 shows a scatterplot for phonological and executive skill factors, where the blue points represent the cases who did not complete the CAT. Overall, these results suggest the two groups were not significantly different from each other.

**Figure 1.**
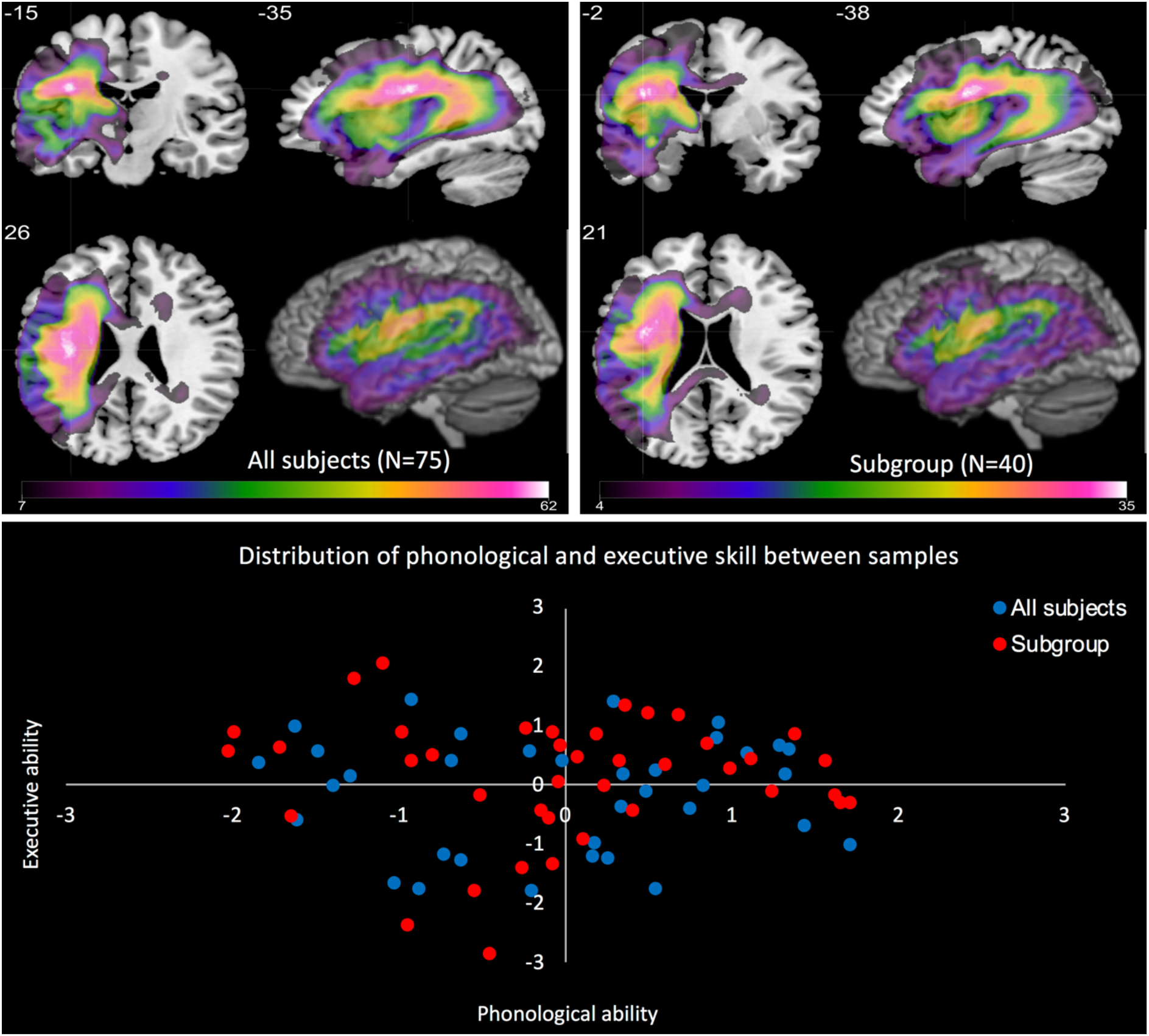
Lesion overlap map for all subjects (top left) and subgroup (top right) in MNI space. The crosshair in both images is located at the maximum lesion overlap. The lower panel shows the distribution of phonological and executive skill component scores for all subjects (blue and red combined) and subgroup (red only).

The remaining results are split into three parts. In the first part, we directly compare behavioural results obtained on the CAT with the extensive test battery. Next, we extract the underlying structure of each battery (extensive, reduced and CAT) and finally, we use the principal component scores from each battery and map them to brain lesions (using both univariate and multivariate models).

### 3.2 Direct comparisons

We compared equivalent or near equivalent tests in the extensive battery and the CAT. We matched seven subtests within the CAT (digit span, repetition of words and non-words, comprehension of spoken words [CSW], comprehension of written words [CWW], semantic memory and object naming) to nine tests from the extensive battery (digit span, PALPA 8 and 9, spoken and written word-to-picture matching, camel and cactus test (pictures), 96-item synonym judgement test, Cambridge naming test, and Boston naming test). All tests have control cut-off scores (obtained from Thompson et al., 2018) except for digit span, PALPA 9 and BNT, which were available in the original test manuals. In Figure 2 we present four pair-wise comparisons as examples (repetition, naming, semantic memory and digit span; all detailed comparisons between tests are shown in Supplementary Materials Section 3). Using the cut-off scores for each test, we derived four quadrants. The bottom left quadrant and top right quadrant contains cases who were impaired or in the normal range in both tests, respectively (thus if the tests were in perfect agreement then all cases would fall into these quadrants). The bottom right quadrant represents cases that scored in the normal range on the CAT but were impaired on the extensive test, whereas cases in the top left quadrant were the opposite (i.e. in the normal range on the extensive test but impaired on the CAT). All scores are represented as percentages. Overall, each CAT subtest and its matched extensive test was found to be correlated though this relationship varied from test to test (R^2^ mean = 0.68, STD = 0.18, range = 0.43 – 0.92), being best for repetition and moderate for semantics. The proportion of patients identified within the normal range by the CAT but impaired on the extensive test was higher (mean = 19.69%, STD = 8.81%, range = 7.5 – 35%) than the reverse (mean = 4.06%, STD = 5.82%, range = 0 – 17.5%) (Wilcoxon rank test p = 0.0016).

**Figure 2.**
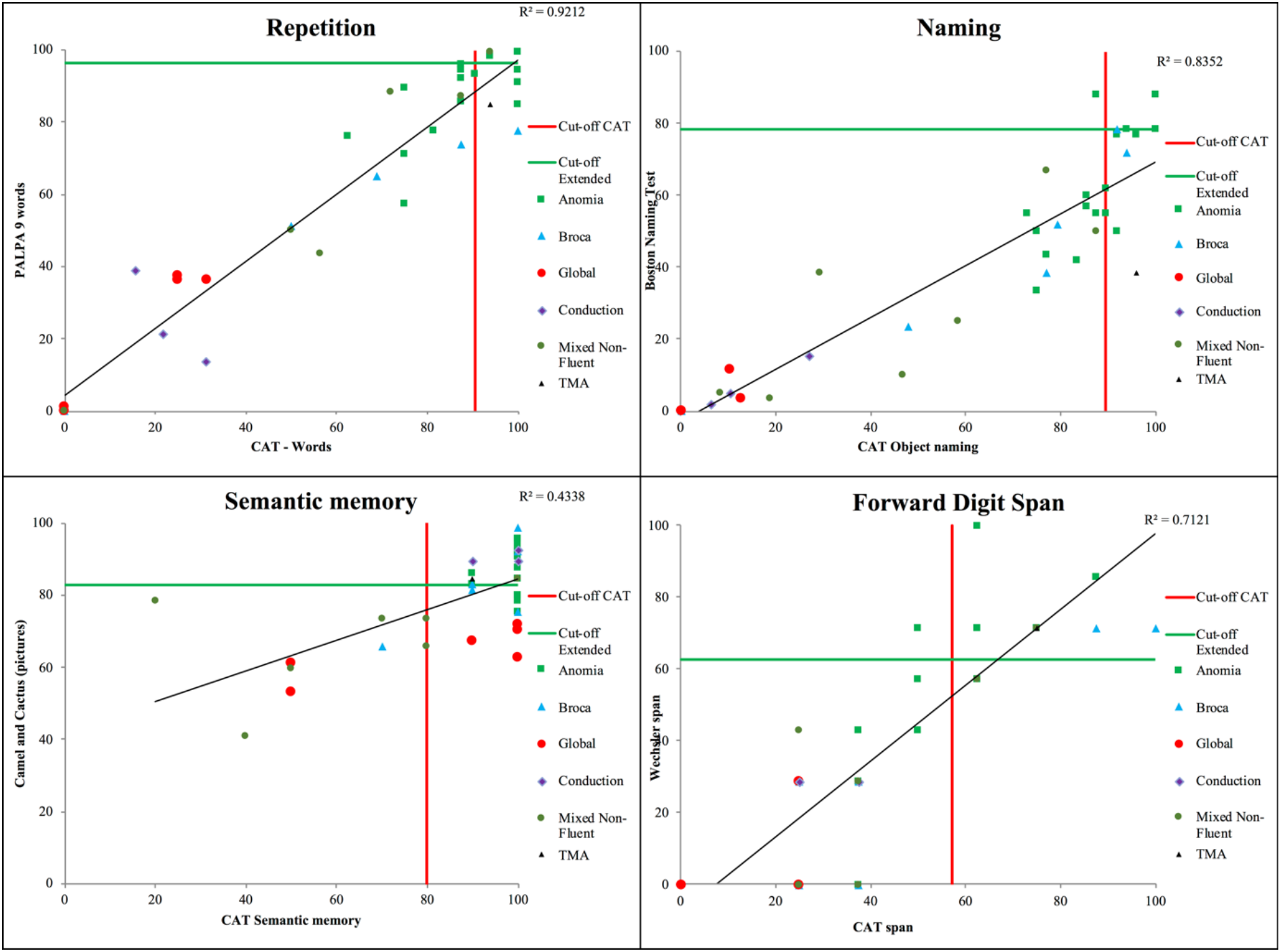
Pairwise comparisons between four example CAT subtests and their matched extensive battery tests: repetition, naming, semantic memory and forward digit span. Each graph has cut-off lines for ‘normal’ performance for the CAT (red line) and extensive (green line) test.

### 3.3 Identifying the underlying structure of language batteries

The k-fold analysis identified a four-factor solution for the extensive and reduced batteries regardless of whether it was computed in the full patient cohort or the subgroup. Only a two-factor solution was identified for the CAT subgroup. Generalisability of the PCA models to the left-out cases was very high for all batteries and cohorts: extensive battery with all cases (r = 0.88), extensive battery with subgroup (r = 0.88), reduced battery with all cases (r = 0.89), reduced battery with subgroup (r = 0.90) and CAT with subgroup (r = 0.79).

Figure 3 shows the factor loadings for each of the PCA solutions. The PCA on the extensive battery with all cases replicated previous findings (Halai *et al.*, 2017, 2018). This PCA model explained 76.7% of the variance and was split into ‘phonological skill’ (accounting for 32.5% variance), ‘executive function’ (16.8% variance), ‘speech quanta’ (13.8% variance) and ‘semantics’ (13.6% variance). These components were replicated in the other iterations of the extensive and reduced test batteries. The extensive battery on the subgroup (77.9% total variance explained) produced the following model: ‘phonological skill’ (34.1% variance), ‘executive function’ (18.9% variance), ‘speech quanta’ (14.3% variance) and ‘semantics’ (10.7% variance). The reduced battery on all cases (78.5% total variance explained) produced the following model: ‘phonological skill’ (28.6% variance), ‘executive function’ (15.1% variance), ‘speech quanta’ (17.5% variance) and ‘semantics’ (17.5% variance). The reduced battery on the subgroup (80.4% total variance) produced the following model: ‘phonological skill’ (29.9% variance), ‘executive function’ (15.0% variance), ‘speech quanta’ (17.6% variance) and ‘semantics’ (18.0% variance). Correlational analyses measuring similarity across these components confirmed very high correlation values between equivalent components (r’s > 0.95, p<0.001), regardless of sample size or battery used, suggesting that the underlying PCA structure obtained on the extensive or reduced battery was stable and equivalent.

**Figure 3.**
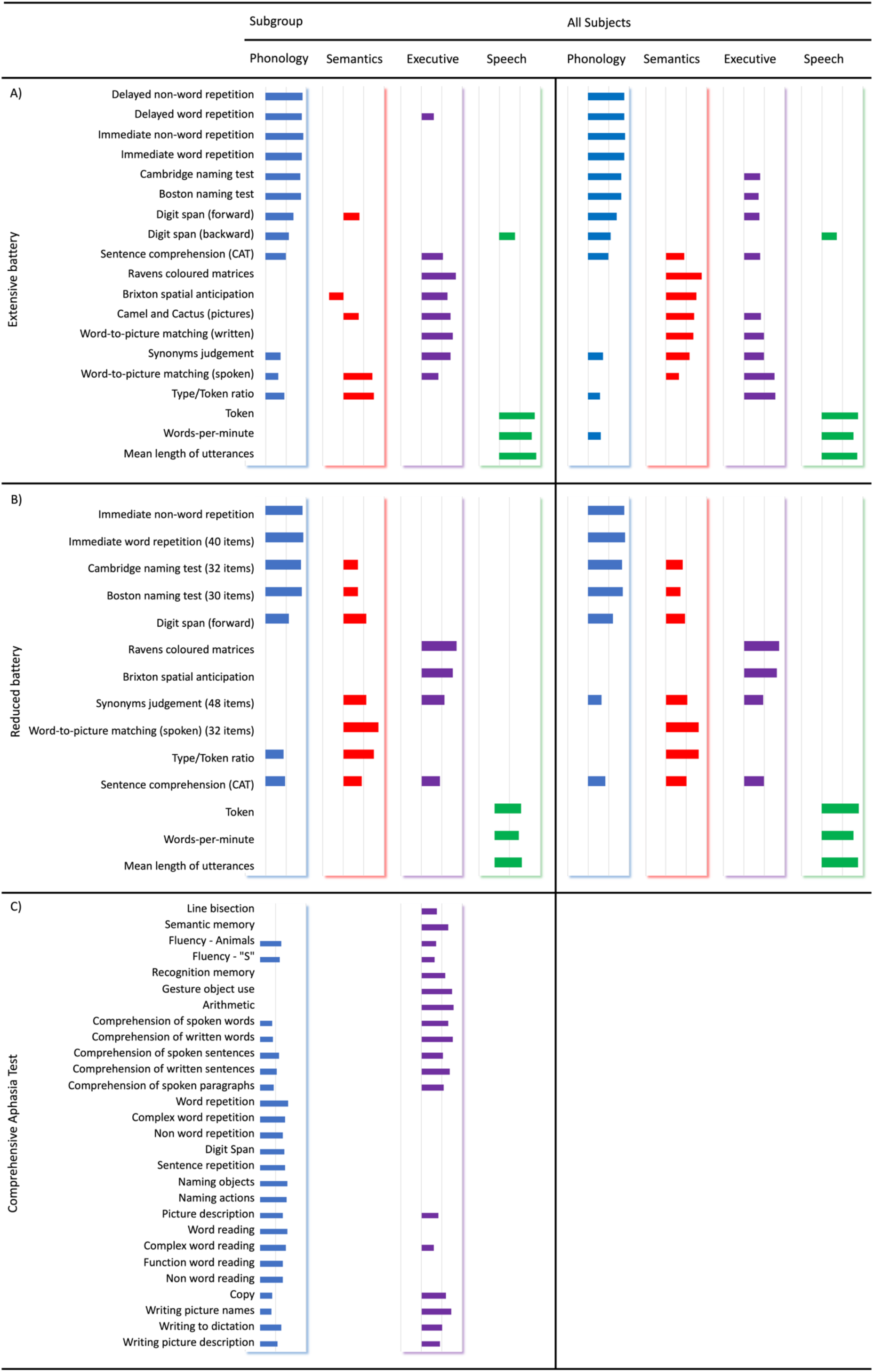
Composite figure showing test loadings for five principal component analyses: a) extensive battery on all cases and subgroup, b) reduced battery on all cases and subgroup, and c) comprehensive aphasia test on subgroup. Loadings between −0.2 – 0.2 are omitted for clarity as they represent weak relationships to the components. The colour coding corresponds to each component: phonology (blue), semantics (red), executive (purple) and speech quanta (green).

The results for the CAT were different. A two-factor solution was obtained with the model explaining 63.1% of the variance. The first factor (accounted for 39.5% variance) was loaded onto by tests requiring speech production and complex comprehension; hence, the factor was termed phonological-language severity. The second factor (23.6% variance) included all other tests not involving phonological production. These tests also varied in difficulty and so we termed this factor overall cognitive severity. Three tests load on both factors (writing to dictation and comprehension of spoken and written sentences). Line bisection did not load onto any factor (nor was it sufficient to create a third component in this data) as it does not measure language or cognitive performance. The interpretation of the two components are supported by finding correlations between the first CAT component with both the phonology and semantics dimensions derived from the full battery (r = 0.88 p < 0.001 and r = 0.44 p < 0.005, respectively) and the second CAT component with the executive ability dimension (r = 0.79, p<0.001).

In summary, a four-factor solution was obtained with the extensive battery. This solution was highly stable and was maintained when the test battery was reduced and when applied to the smaller patient subgroup. In contrast, the CAT data produced a two-factor solution, where the first component related to phonological-language severity and the second component was related to executive or generalised cognitive severity.

### 3.4 Mapping brain-behaviour relationships

The univariate results are summarised in Figure 4, which shows significant clusters for every principal component across different behavioural batteries (detailed peak co-ordinate information is provided in Supplementary Materials Section 4). All results were conducted with and without lesion volume correction but, for brevity, we only present the lesion corrected results here (uncorrected results are in Supplementary Materials Section 5). The neural correlates replicated previous findings (Halai et al., 2017): 1) phonology was related to the integrity of the superior temporal gyrus extending posteriorly into supramarginal gyrus and angular gyrus; 2) semantic ability related to the integrity of the middle and ventral temporal cortex extending posteriorly into occipital cortex; and 3) speech quanta was related to precentral gyrus and inferior frontal gyrus. We extended previous findings by identifying a large posterior cluster for executive skill centred on the lateral occipital cortex. This result was replicated across the full vs. data-driven reduced batteries and in the full cohort vs. patient subgroup. The two univariate neural correlates of the CAT components highly overlapped with the phonological and executive clusters from the extensive battery (in keeping with the behavioural correlations noted above). We compared results across batteries to determine if there were significant differences in their statistical maps; all unthresholded t-maps were converted into z-maps (using SPM12 function spm_t2z.m) and pair-wise difference maps were obtained for equivalent components. We did not find any differences for any comparison (z threshold ±3.29 and arbitrary cluster extent > 100).

**Figure 4.**
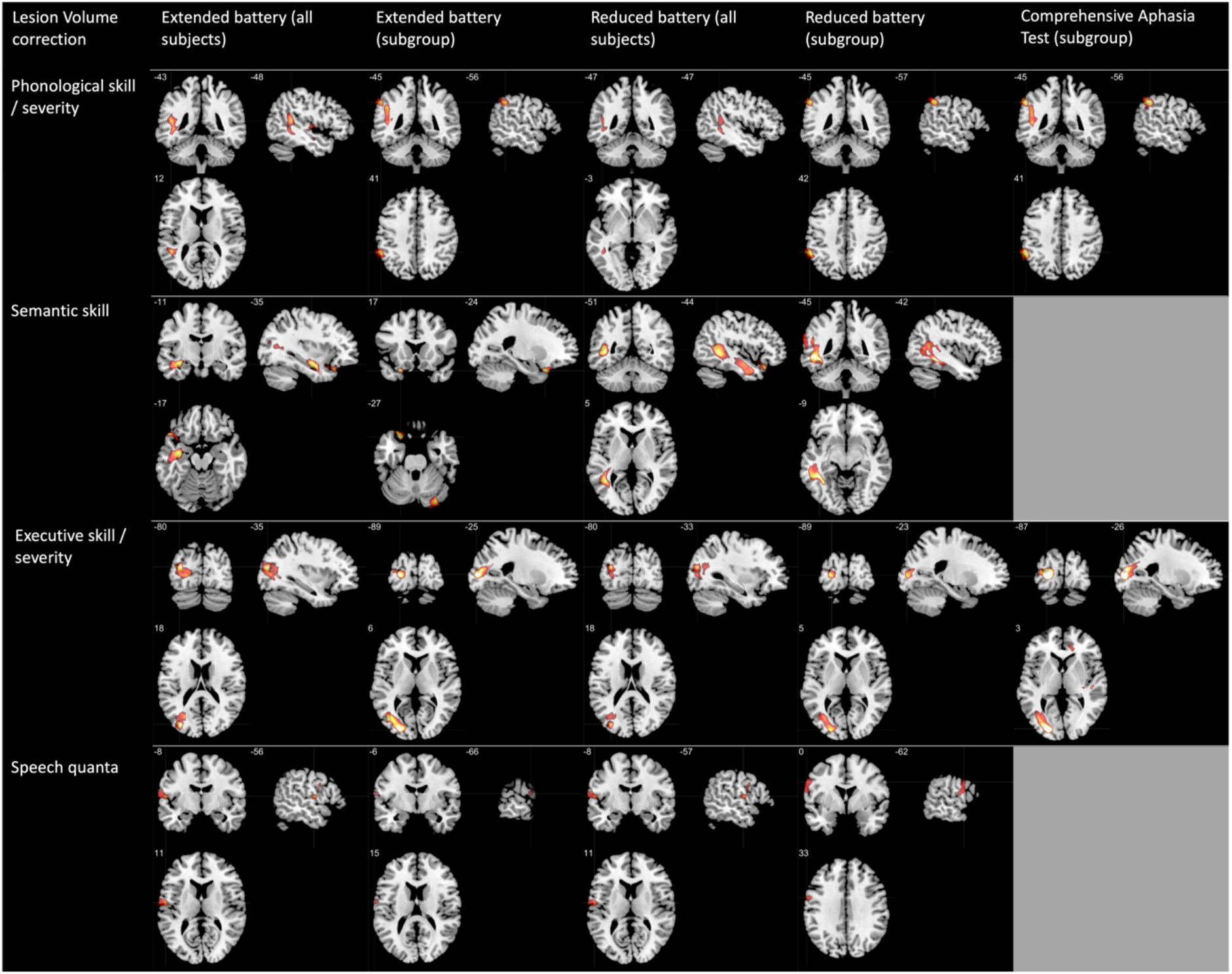
VBCM results for all components with lesion volume correction using voxelwise p<0.001 and family wise error cluster correction p<0.05 (except the speech quanta cluster for the extensive battery [subgroup], which is thresholded using a voxelwise p < 0.002 and family wise error cluster correction p < 0.05). The rows represent each principal component; phonological skill / severity, semantic skill, executive skill/severity and speech quanta. The grey patches in the final column indicate that there were no corresponding CAT components for semantic skill and speech quanta. Each panel has a cross hair located at the peak voxel. Scale t-values = 3 - 5.

Briefly, we note that the results when lesion volume correction was omitted were almost identical to those stated above. As expected, cluster sizes were larger without lesion volume correction but their locations were generally convergent with the clusters found with lesion volume correction. There was one minor exception: the speech quanta cluster in the extensive battery for the subgroup was significant at the typical threshold of p<0.001 voxelwise, with FWEc corrected p<0.05 (as opposed to p<0.005 voxelwise, with FWEc corrected p<0.05 with lesion volume correction).

Finally, we present results from the multivariate analyses. Table 1 shows the cross-validated correlation coefficients and corresponding p-values for each model. Each analysis was performed twice, with lesion volume either included or excluded as a covariate. For models without lesion volume correction, the phonological skill component was predicted for the extensive battery (all and subgroup) (cross validated r = 0.30 and 0.35, respectively) and the equivalent CAT component (cross validated r = 0.37). Results were not significant for the phonological skill component obtained from the reduced battery on the subgroup but was at trend with all cases (cross validated r = 0.22, p = 0.075). In contrast, the semantic component was successfully predicted in all batteries (cross validated r’s > 0.43) apart from the CAT (which produces no equivalent principal component). The executive component was predicted for all batteries, including the equivalent CAT component (cross validated r’s > 0.30). The speech quanta component was only predicted in the extensive and reduced battery for all cases (cross validated r = 0.25 and 0.26, respectively). For models with lesion volume correction, models significantly predicting semantic component scores were obtained for all batteries (cross validated r’s > 0.27), again apart from the CAT (which produces no equivalent principal component). The executive component was predicted for the extensive and reduced subgroups (cross validated r = 0.49 and 0.36, respectively), while the same batteries for all cases were at trend (cross validated r’s > 0.19, p’s < 0.096). Finally, three models for phonological skill were at trend, including the extensive and reduced batteries for all cases and the reduced battery for the subgroup (cross validated r’s > 0.19, p’s < 0.096). The remaining models were not significant. In summary, the results for the extensive battery on all cases was very similar to the reduced battery for all cases (with and without lesion volume correction). The speech quanta component is poorly predicted in the subgroups for the extensive and reduced batteries (with and without lesion correction). Finally, CAT component scores were not significant predicted by multivariate lesion modelling when lesion volume correction is applied.

## Discussion

Cognitive and language deficits due to brain injury or progressive disorders are typically multifaceted and can range from severe to very mild symptoms. There is a pressing need to be able to detect neuropsychological deficits across a wide range of severities and domains in a time frame that is feasible for application in clinical and research settings. Most clinical test batteries approach this problem by adopting a “shallow” battery that tests a wide range of deficits using a small number of trials (typically < 10 per domain tested). Shallow batteries can generate a quick impression of patients’ strengths and weaknesses across many different domains, which can be followed up with more detailed, targeted assessment. The limited dynamic range in each assessment, however, can be problematic for core clinical and research needs. Specifically, short subtests can be insensitive to mild impairments, struggle to grade different levels of impairment, and fail to detect longitudinal change. Such limitations are problematic in the clinic and research (e.g., missing mild impairments, inability to detect changing performance, insufficient test score variance for correlation-based analyses such as lesion symptom mapping). This potential inability to seriate patients is also potentially problematic for investigating clinical disorders, such as post-stroke aphasia (PSA) that exhibit graded variation along continuous behavioural dimensions (Butler *et al.*, 2014; Corbetta *et al.*, 2015, Mirman *et al.*, 2015*a*; Lacey *et al.*, 2017; Halai *et al.*, 2018; Schumacher *et al.*, 2019). To explore these important clinical and research issues, the current study used the test case of PSA where there is a long history of using systematic multi-domain test batteries. Specifically, we compared an extensive, detailed test battery against a “shallow” assessment battery (the Comprehensive Aphasia Test; CAT) and then generated a new, data-driven battery which preserved the depth but reduced the number of tasks. For all three batteries we explored their ability to reveal the graded, multidimensional structure that underpins PSA and also their lesion correlates.

Overall, our results show that multiple subtests in the CAT were less sensitive to mild impairments than in the extensive battery (on average 19.69% cases missed) and the correlations between the tests, whilst good in general (average R^2^ = 0.68), varied (being best for repetition and weakest for semantics). Indeed, semantic deficits were harder to detect in the CAT, with 30-35% of impaired cases missed depending on the task. Cross-validated PCA of the extensive battery showed that there were four, very robust dimensions of variation (phonology, semantics, fluency and cognitive-executive skill). In contrast, the CAT only generated two dimensions (phonology-language and generalised cognition) which, in the case of language spanned two of the components derived from the full battery. We successfully used PCA to derive a new reduced battery that allowed a data-drive reduction in the number of tests and also reduced the number of items in some of the longer assessments. As intended, this reduced battery retained the four, robust language and cognitive components. Finally, in a series of univariate and multivariate lesion-symptom mapping analyses, the same pattern of results emerged; the full and data-driven reduced batteries revealed the same discrete areas associated with each of the four PCA components, whilst the CAT generated two areas of interest that overlapped with a subset of those observed from the alternative batteries.

It is, of course, important to consider the targets of investigation before selecting the most suitable assessments. The psycholinguistically-informed CAT was designed to provide a broad sampling of many different language activities through a ‘shallow’ test design. This is the common approach to saving assessment time though, as demonstrated in the current study, it is also possible to use an alternative approach in which time is saved by reducing the number of tests but preserving the depth of each test. The latter approach, by definition, cannot sample many different activities but the greater number of test items allow it to be sensitive to mild impairments and grade impairments. The resulting larger dynamic range can be important in both the clinical and research; for example when needing to measure change over time (e.g., to track decline in progressive disorders, performance improvements in spontaneous recovery or after intervention, etc.) or when relating variation in language-cognitive performance to other factors and the distribution of underlying brain damage. The ability to fathom the underlying behavioural variations using PCA is also very likely to reflect the available dynamic range in the tests (like any correlation-based analysis, PCA requires sufficient variation to be present). Whilst it can be important to assess performance on specific activities, the PCA results from this large and diverse PSA cohort indicate that a large proportion of the total cohort variation (~80%) can be captured by four orthogonal dimensions. This follows from the facts that (a) each task is not “pure” but instead reflects a combination of core language and cognitive skills and (b) that, resultantly, there is considerable collinearity across different tests (Patterson and Lambon Ralph, 1999; Butler *et al.*, 2014; Halai *et al.*, 2017). PCA also provides a data-driven solution to the question; which subset of tests should be selected from an extensive battery? The same multidimensional variation can be captured by selecting a subset of tasks that are aligned with only one of the principal components.

Finally, we discuss the neural correlates and multivariate prediction results for the components scores across the different test batteries. The univariate VBCM analysis identified separable neural correlates for all component scores across all test batteries. The clusters were highly convergent with recent reports that have found: 1) phonology to be related to the supramarginal gyrus but extending into posterior superior temporal gyrus (Hickok and Poeppel, 2007; Price, 2012; Butler *et al.*, 2014; Halai *et al.*, 2017, 2018); 2) semantics to be related to anterior inferior and middle temporal gyrus (Lambon Ralph *et al.*, 2017); and 3) speech quanta being related to precentral gyrus extending into the insula (Borovsky *et al.*, 2007; Kinoshita *et al.*, 2015; Halai *et al.*, 2017). The current study also identified regions in the left occipital, posterior temporal and posterior parietal lobe that were related to executive ability. There is evidence that the lateral temporo-occipital areas are activated for demanding visuo-spatial tasks (Fedorenko *et al.*, 2013; Humphreys and Lambon Ralph, 2017) or when location and feature information must be combined (Simpson *et al.*, 2011). These processes are required when completing the Raven’s Coloured Progressive Matrices and Brixton Spatial Anticipation Test, which loaded highly with the executive component. Other recent investigations of the PSA population have found that executive ability is correlated with superior frontal and paracingulate regions (Geranmayeh *et al.*, 2017; Lacey *et al.*, 2017; Alyahya *et al.*, 2018; Schumacher *et al.*, 2019). One explanation for the discrepancy might relate to the pattern of middle cerebral artery (MCA) lesions observed in a typical stroke population, whereby the highest probability of damage occurs in the striatocapsular region and insula (Phan *et al.*, 2005) and only very large MCA strokes damage the superior frontal and occipital regions (as they fall in watershed regions of the anterior cerebral and posterior cerebral artery, respectively). This would support the generally accepted hypothesis that increased lesion size is consistent with increased behavioural deficits, both language and executive.

Interestingly, the pattern of neural correlates across the components within different test batteries was remarkably similar. This probably reflects the fact that the batteries seem to assess the same four underlying dimensions. Even for the CAT, the lesion correlates for its two PCA components were almost identical to the clusters found for phonology and executive skills in the extensive battery. The ability to predict the component scores using lesion information was also highly consistent when all cases were used in the extensive and reduced battery. The lesion data was able to predict all components without lesion volume correction and 3/4 tests with lesion volume correction (although some models were at trend). Results were mixed for the subgroup batteries, such that the models typically failed at predicting phonology and speech quanta. One reason for the lack of consistency might simply be due to the sample size, since multivariate decoding methodologies typically require large samples as data are partitioned into train/test sets for cross-validation. A recent simulation study (Sperber *et al.*, 2019) suggested that approximately 100 subjects are required to have stable/reproducible beta parameter mapping, whereas for prediction of clinical outcomes the number peaked at 40 and was relatively stable from this point up to 100 cases. The numbers in the current study reflect these two ranges: 75 for the extensive battery (which generated robust results) and 40 for the subgroup analyses.

## Abbreviations

CAT: Comprehensive Aphasia Test
BDAE: Boston Diagnostic Aphasia Examination
WAB: Western Aphasia Battery
MTDDA: Minnesota Test for Differential Diagnosis of Aphasia
PICA: Porch Index of Communicative Ability
PSA: Post Stroke Aphasia
PCA: Principle Component Analysis
PALPA: Psycholinguistic Assessment of Language Processing in Aphasia
CCTp: Camel and Cactus pictures
BNT: Boston Naming Test
T: Tokens
WPM: Words Per Minute
MLU: Mean Length of Utterances
TTR: Type Token Ratio
VBCM: Voxel Based Correlational Methodology
PRoNTo: Pattern Recognition of Neuroimaging Toolbox
FWEc: Family Wise Error corrected
CSW: Comprehension Spoken Words
CWW: Comprehension Written Words
MCA: Middle Cerebral Artery

## Acknowledgements

We thank all the patients, families, carers and community support groups for their continued, enthusiastic support of our research programme.

## Supplementary Materials

### Section 1

**Table 1.** Demographic information for the full sample of cases with chronic post stroke aphasia and the subgroup. Cases are ordered by lesion volume. Abbreviations: Transcortical sensory aphasia (TSA); Transcortical motor aphasia (TMA)

| ID | Gender | Aphasia<br>classification | Age | Years of<br>education | Months<br>post-stroke | Lesion<br>volume | Sub-<br>group |
| --- | --- | --- | --- | --- | --- | --- | --- |
| 1 | F | Anomia | 43 | 16 | 15 | 175 | * |
| 2 | M | Anomia | 75 | 11 | 12 | 1481 |  |
| 3 | F | Anomia | 53 | 11 | 47 | 1526 |  |
| 4 | M | Anomia | 68 | 11 | 21 | 3311 | * |
| 5 | M | Broca | 61 | 11 | 16 | 3528 |  |
| 6 | F | Conduction | 46 | 16 | 21 | 3897 | * |
| 7 | M | Anomia | 50 | 19 | 16 | 4538 | * |
| 8 | M | Conduction | 68 | 11 | 37 | 4773 |  |
| 9 | M | Anomia | 65 | 17 | 25 | 4806 |  |
| 10 | M | Conduction | 67 | 11 | 13 | 4879 | * |
| 11 | F | Broca | 48 | 12 | 16 | 5273 |  |
| 12 | M | TSA | 63 | 12 | 24 | 5822 |  |
| 13 | M | Conduction | 67 | 17 | 14 | 6557 | * |
| 14 | M | Anomia | 65 | 10 | 85 | 6607 | * |
| 15 | F | Wernicke | 77 | 16 | 34 | 6843 |  |
| 16 | M | Anomia | 56 | 16 | 17 | 6974 | * |
| 17 | F | Anomia | 51 | 11 | 66 | 6975 | * |
| 18 | M | Anomia | 84 | 9 | 35 | 7854 |  |
| 19 | F | Anomia | 68 | 16 | 22 | 8118 |  |
| 20 | M | Anomia | 87 | 12 | 35 | 8238 |  |
| 21 | M | Anomia | 44 | 11 | 40 | 8437 | * |
| 22 | M | Anomia | 86 | 9 | 17 | 8528 | * |
| 23 | M | Mixed Non-fluent | 68 | 11 | 14 | 8788 | * |
| 24 | F | Anomia | 73 | 11 | 89 | 8921 |  |
| 25 | F | Anomia | 69 | 19 | 39 | 9159 | * |
| 26 | F | Mixed Non-fluent | 77 | 11 | 20 | 9229 | * |
| 27 | F | Anomia | 52 | 12 | 76 | 9767 | * |
| 28 | F | Mixed Non-fluent | 51 | 11 | 40 | 10051 | * |
| 29 | M | Anomia | 67 | 11 | 60 | 10073 |  |
| 30 | M | TMA | 76 | 11 | 116 | 11239 |  |
| 31 | M | Broca | 85 | 10 | 46 | 11393 |  |
| 32 | M | Broca | 52 | 17 | 33 | 11915 | * |
| 33 | F | Mixed Non-fluent | 75 | 11 | 160 | 12057 | * |
| 34 | M | Broca | 82 | 10 | 13 | 12131 | * |
| 35 | F | Anomia | 58 | 11 | 278 | 12699 | * |
| 36 | M | Broca | 59 | 13 | 37 | 13080 | * |
| 37 | M | Global | 78 | 11 | 17 | 13187 | * |
| 38 | F | Anomia | 77 | 11 | 56 | 13577 | * |
| 39 | M | Mixed Non-fluent | 58 | 13 | 32 | 14625 |  |
| 40 | M | Anomia | 56 | 11 | 26 | 14681 | * |
| 41 | M | Global | 66 | 11 | 12 | 14890 |  |
| 42 | M | Anomia | 66 | 11 | 126 | 15492 |  |
| 43 | M | Anomia | 80 | 11 | 84 | 15857 | * |
| 44 | M | Anomia | 59 | 11 | 34 | 16433 | * |
| 45 | M | Broca | 80 | 12 | 65 | 18163 |  |
| 46 | M | Broca | 58 | 11 | 135 | 18392 | * |
| 47 | M | Broca | 54 | 13 | 35 | 18632 | * |
| 48 | M | Anomia | 63 | 12 | 12 | 18639 |  |
| 49 | F | Anomia | 44 | 13 | 37 | 18948 | * |
| 50 | M | Global | 74 | 11 | 18 | 19500 |  |
| 51 | M | Broca | 51 | 12 | 34 | 20043 |  |
| 52 | M | Anomia | 85 | 10 | 69 | 21489 |  |
| 53 | M | Mixed Non-fluent | 73 | 11 | 23 | 22732 | * |
| 54 | M | Anomia | 51 | 13 | 72 | 22948 | * |
| 55 | F | TMA | 73 | 11 | 46 | 23863 | * |
| 56 | M | Mixed Non-fluent | 67 | 11 | 120 | 26097 |  |
| 57 | M | Broca | 50 | 12 | 16 | 26218 |  |
| 58 | F | Mixed Non-fluent | 67 | 14 | 176 | 26283 |  |
| 59 | F | Broca | 66 | 11 | 63 | 26491 | * |
| 60 | M | Global | 72 | 11 | 42 | 27054 |  |
| 61 | M | Broca | 62 | 11 | 104 | 27242 |  |
| 62 | M | Mixed Non-fluent | 81 | 11 | 69 | 28144 |  |
| 63 | M | Mixed Non-fluent | 67 | 11 | 44 | 31317 |  |
| 64 | M | Mixed Non-fluent | 63 | 12 | 42 | 31599 |  |
| 65 | M | Global | 72 | 11 | 155 | 32981 |  |
| 66 | M | Global | 58 | 13 | 57 | 33239 | * |
| 67 | M | Mixed Non-fluent | 64 | 11 | 29 | 33239 | * |
| 68 | M | Mixed Non-fluent | 79 | 11 | 63 | 33678 |  |
| **69** | M | Mixed Non-fluent | 78 | 13 | 36 | 34242 | * |
| **70** | M | Broca | 73 | 11 | 114 | 36877 | * |
| **71** | M | Global | 52 | 11 | 73 | 37822 | * |
| **72** | M | Mixed Non-fluent | 70 | 11 | 38 | 37850 | * |
| **73** | F | Mixed Non-fluent | 52 | 11 | 99 | 40313 |  |
| **74** | M | Global | 68 | 12 | 50 | 41379 | * |
| **75** | M | Mixed Non-fluent | 76 | 11 | 192 | 42568 |  |

### Section 2

In the following section we show the results of the factor analyses (unrotated single dimension) performed on the individual item level scores of each test (or their condition manipulations). This identified each items’ loading onto the factor explaining the largest amount of variance in the data; the top 50% of items were included. Table 2 shows the items that were included in the reduced tests for: 1) PALPA 9 (word repetition), 2) Boston naming test (BNT), 3) Cambridge semantic battery 64-item picture naming, and 4) 96-item synonym judgement test. Each column in Table 2 shows the top 50% loading items following the factor analysis, with the bottom section showing descriptive statistics of the loading values.

The PALPA 9 test for word repetition consists of 80 items and a one factor solution explained 48.87% of the variance. The BNT has 60 items and a one factor solution explained 39.39% of the variance. As the same Cambridge semantic battery items were used in both the picture naming and word-picture matching tests, we wanted to ensure item consistency across tests. Picture naming had a larger variance of scores in the stroke cohort compared to the spoken word to picture matching test (SD = 33.9 and 11.5, respectively) and so we performed the factor analysis on the Cambridge naming test (CNT). The battery consists of 64 items with two animacy groups (living and non-living). A factor analysis on each dimension showed that the model for the living category explained 44.05% variance and the non-living model explained 46.74% variance. The same reduced item list derived from the CNT test data was used in the reduced spoken word-to-picture matching test. The 96-synonym judgement test is split into six groups along high/low frequency (HF/LF) and high/mid/low imageability (HI/MI/LI) dimensions. A factor analysis on the item scores within each group produced the following models: HF HI 33.50% variance explained; HF MI 26.34% variance explained; HF LI 16.70% variance explained; LF HI 41.80% variance explained; LF MI 24.99% variance explained; LF LI 14.76% variance explained.

**Table 2.**
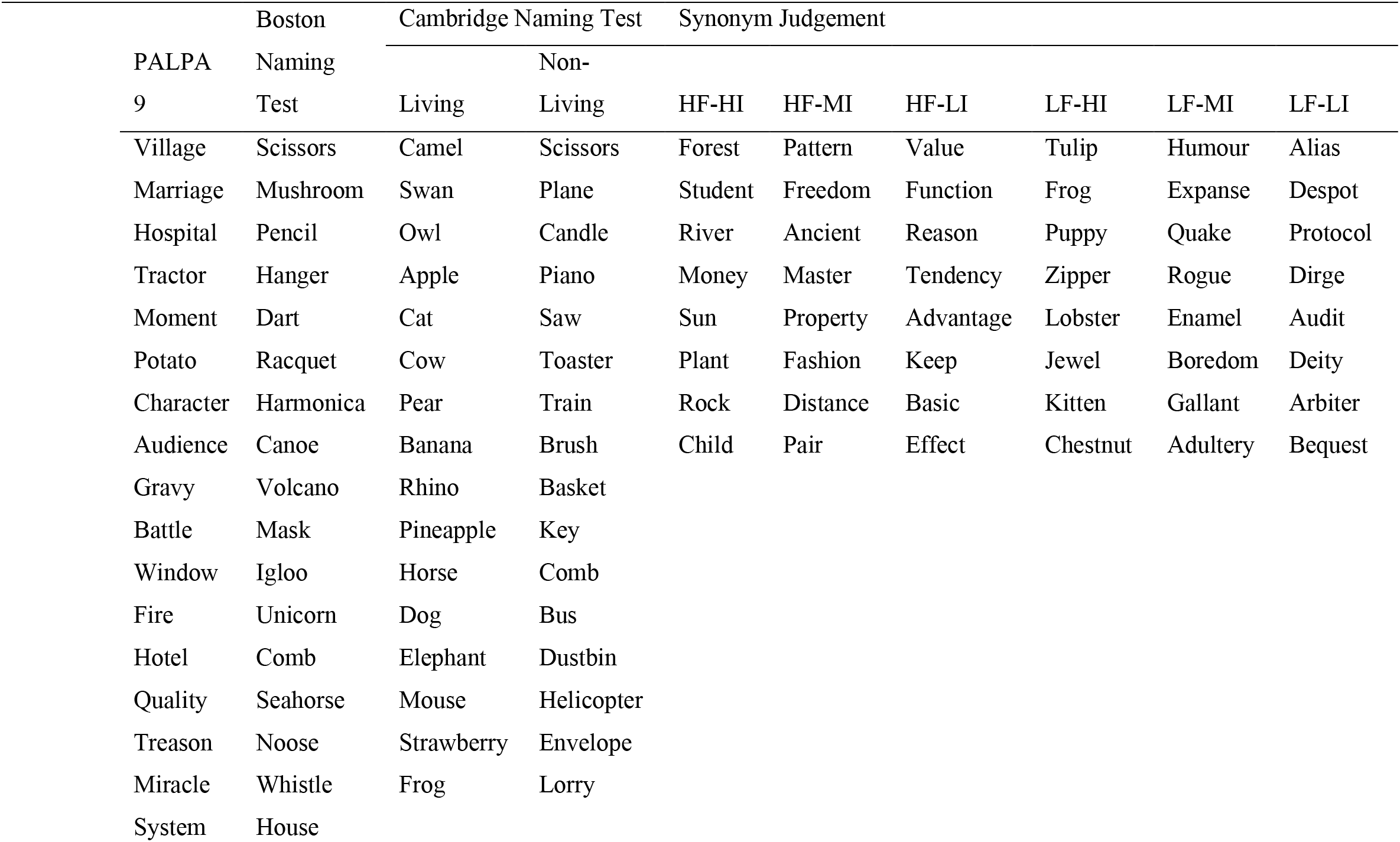
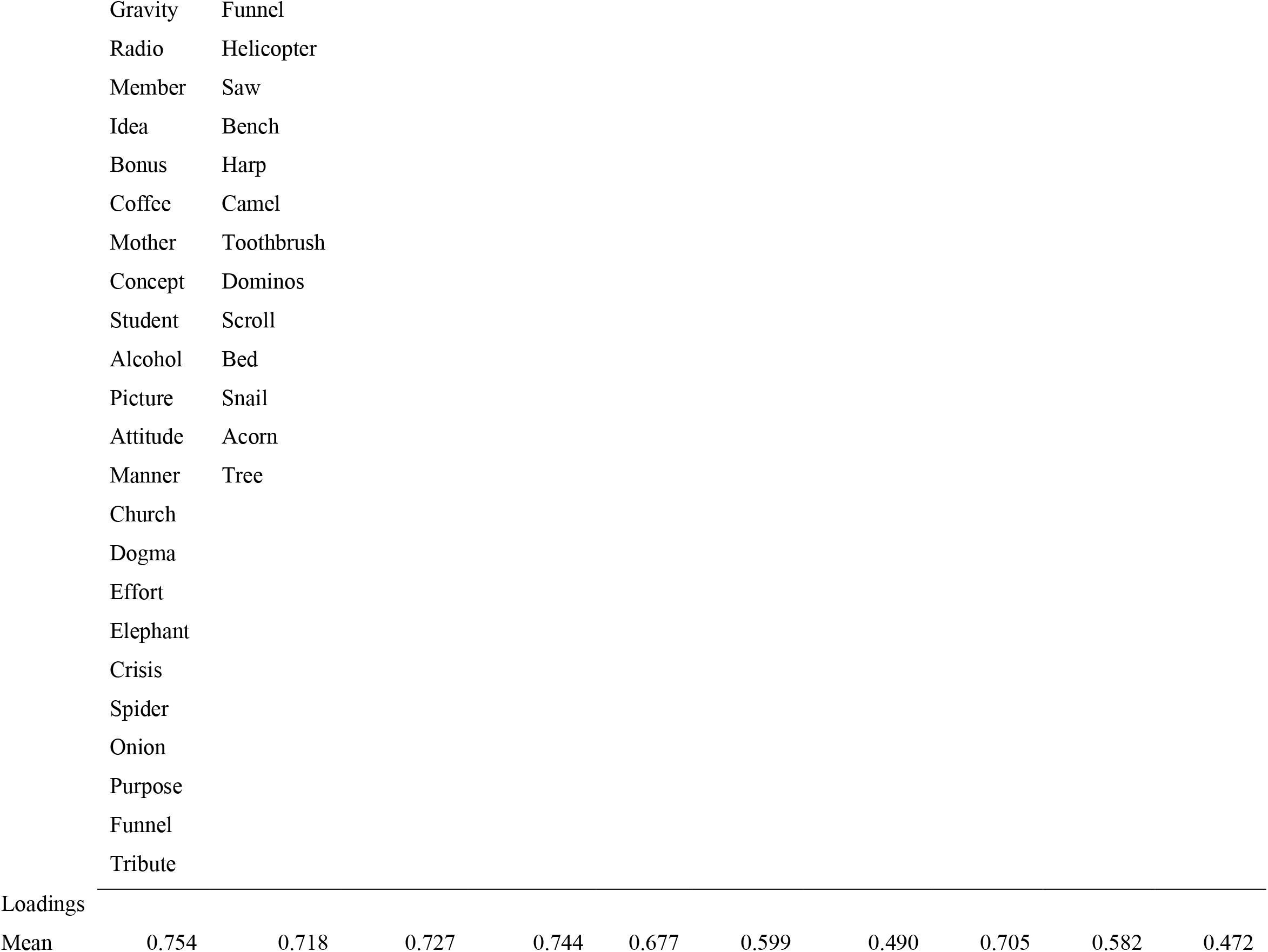
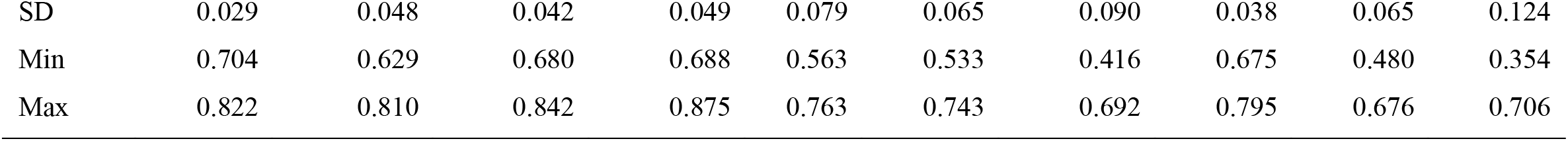
Item list for the reduced tests: 1) psycholinguistic assessment of language processing in aphasia - word repetition (PALPA 9), 2) Boston naming, 3) Cambridge semantic battery 64-item picture naming, and 4) Synonym judgement. Each column is the result of a separate factor analysis and descriptive statistics for the loading of the items are shown at the bottom. Abbreviations: High frequency (HF), low frequency (LF), high imageability (HI), mid imageability (MI), low imageability (LI).

### Section 3

**Table 3.**
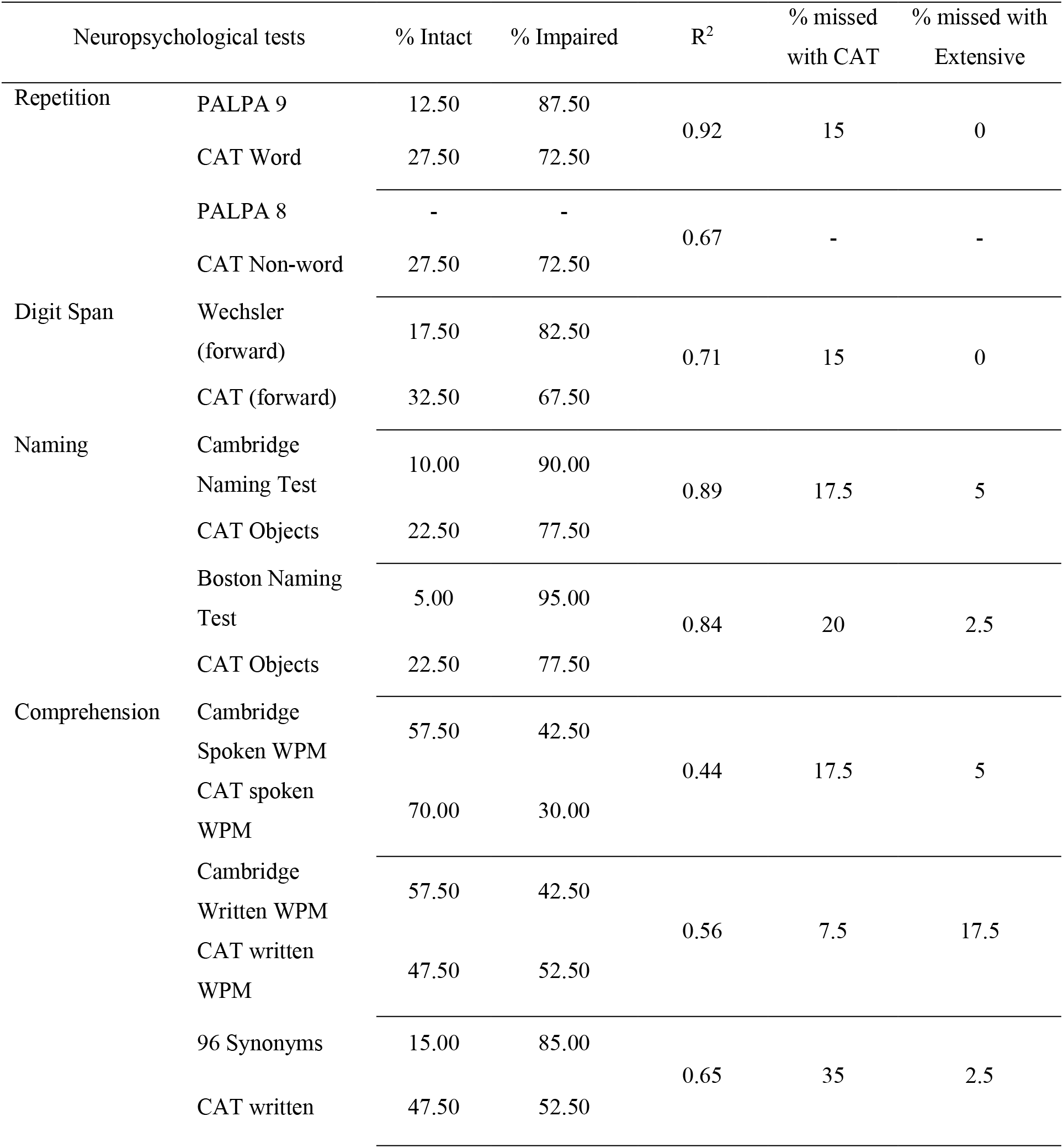
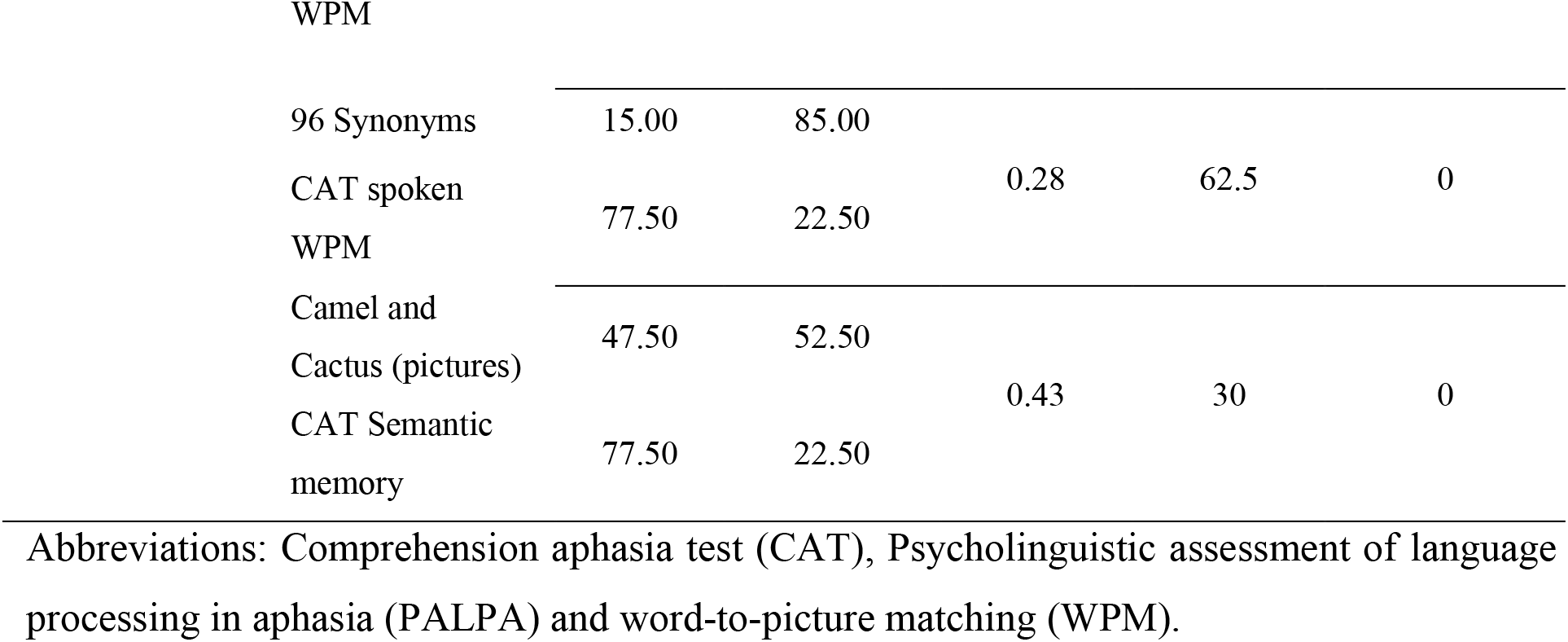
Direct comparison between pair-wise CAT sub-tests and the equivalent extensive test. For each comparison, we show the R^2^ (variance explained) based on correlations, proportion of cases who were determined to have intact or impaired scores (compared to pre-existing norm data taken from the original test batteries or, where none were available, from Thompson et al., 2018). The final two columns indicate the proportion of patients who were identified as impaired on the extensive tests but not on the CAT (% missed with CAT) and those identified as impaired on the CAT but not on the extensive test (% missed with Extensive).

### Section 4

**Table 4.**
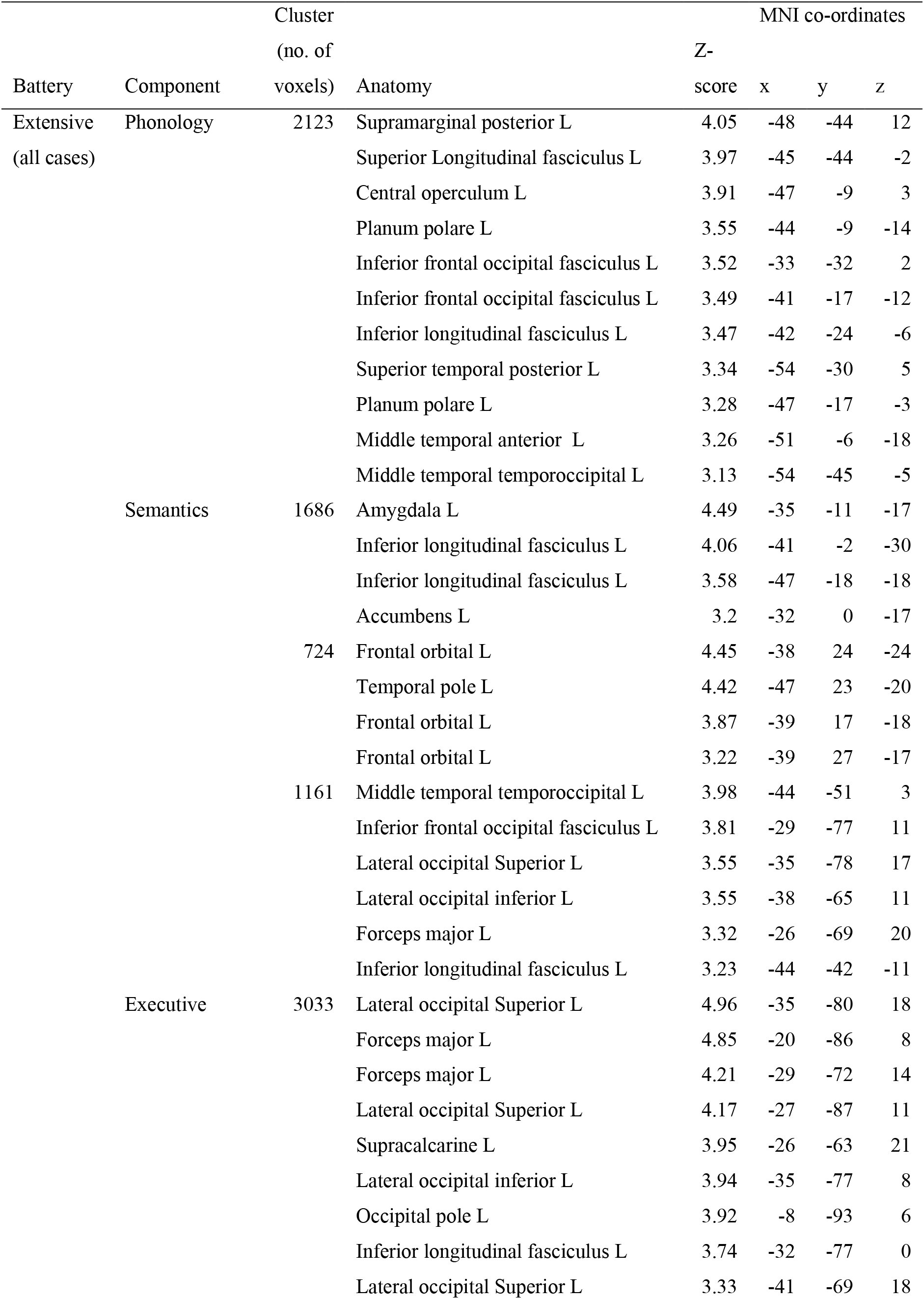
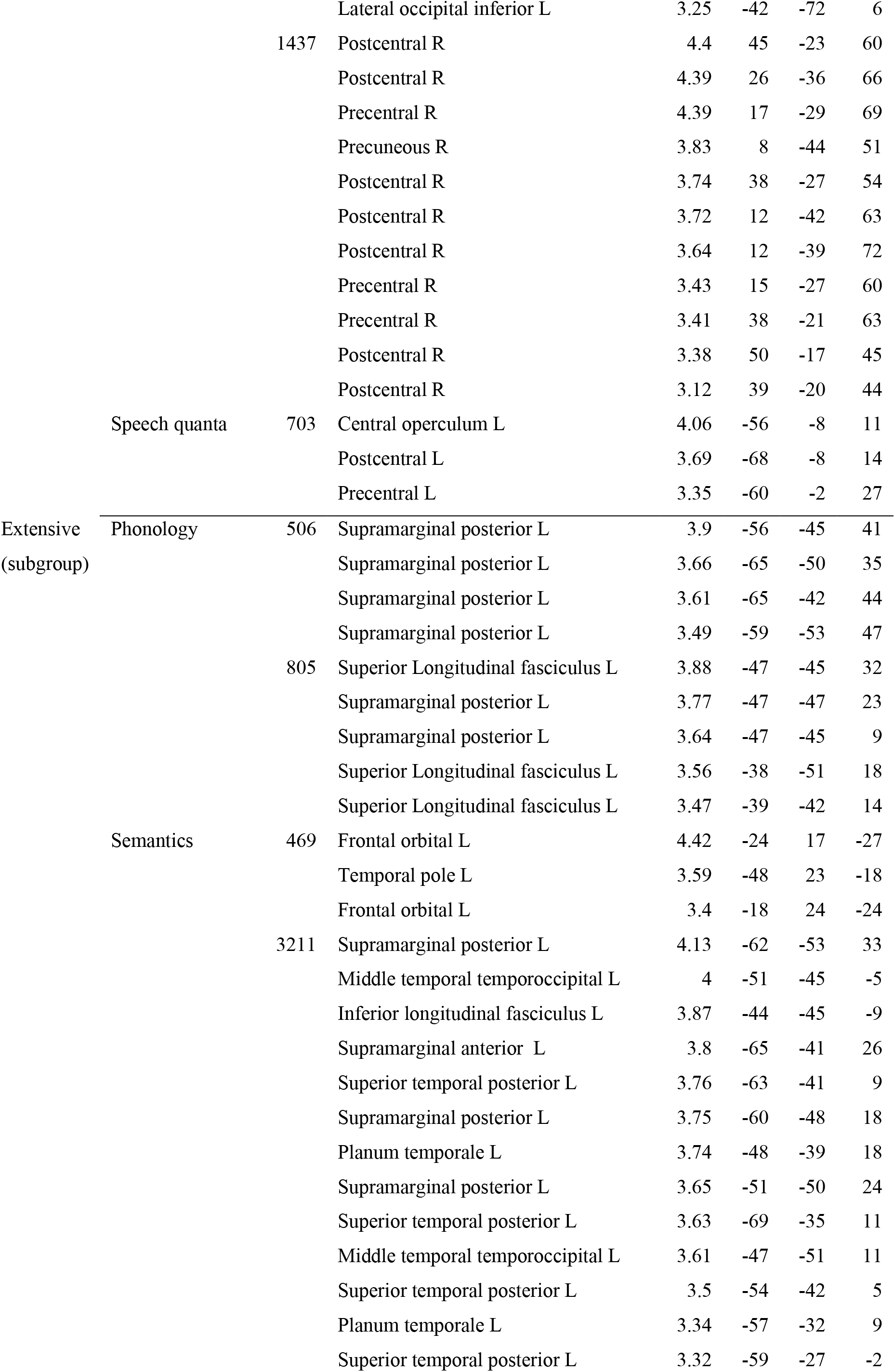
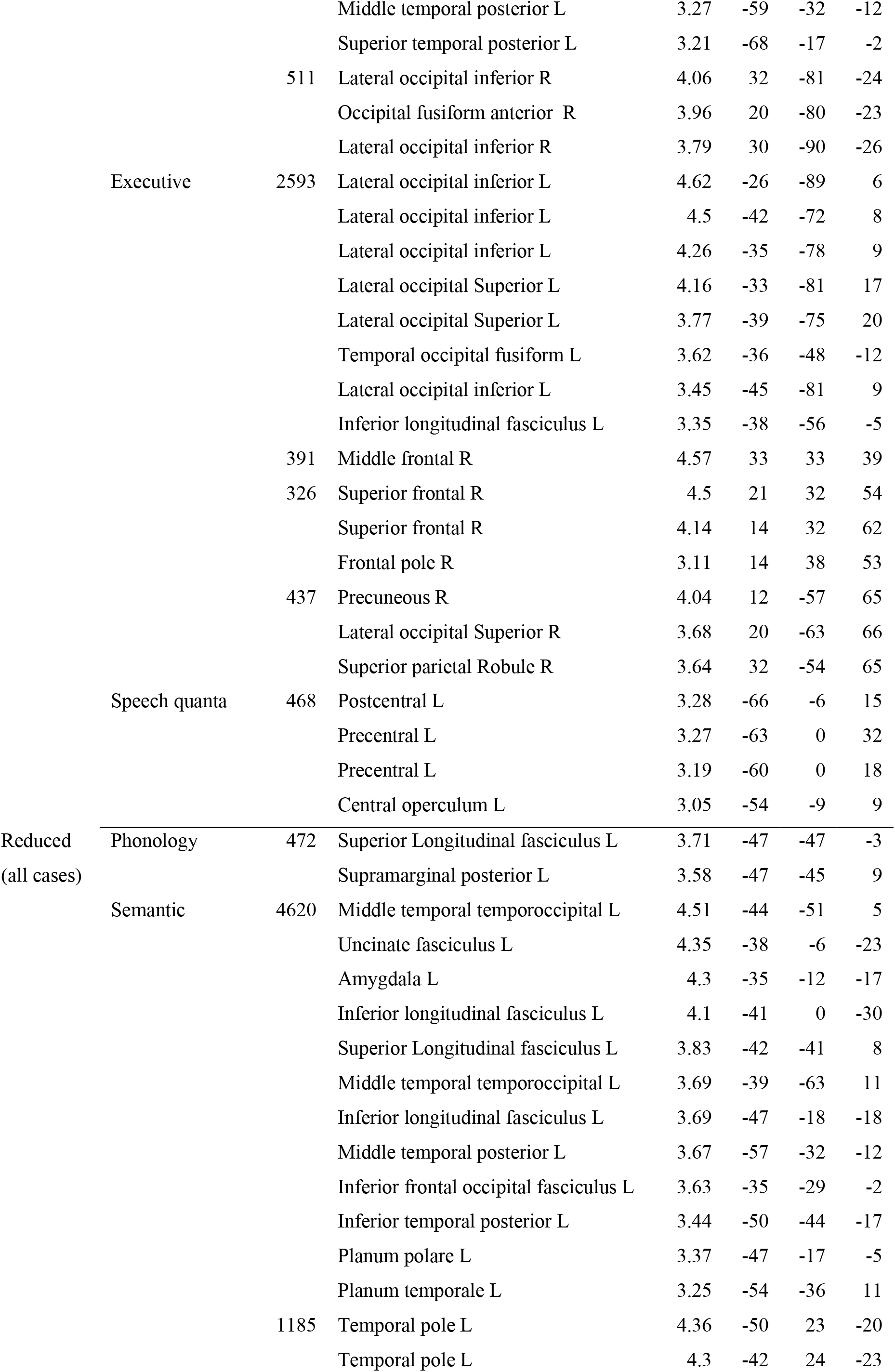
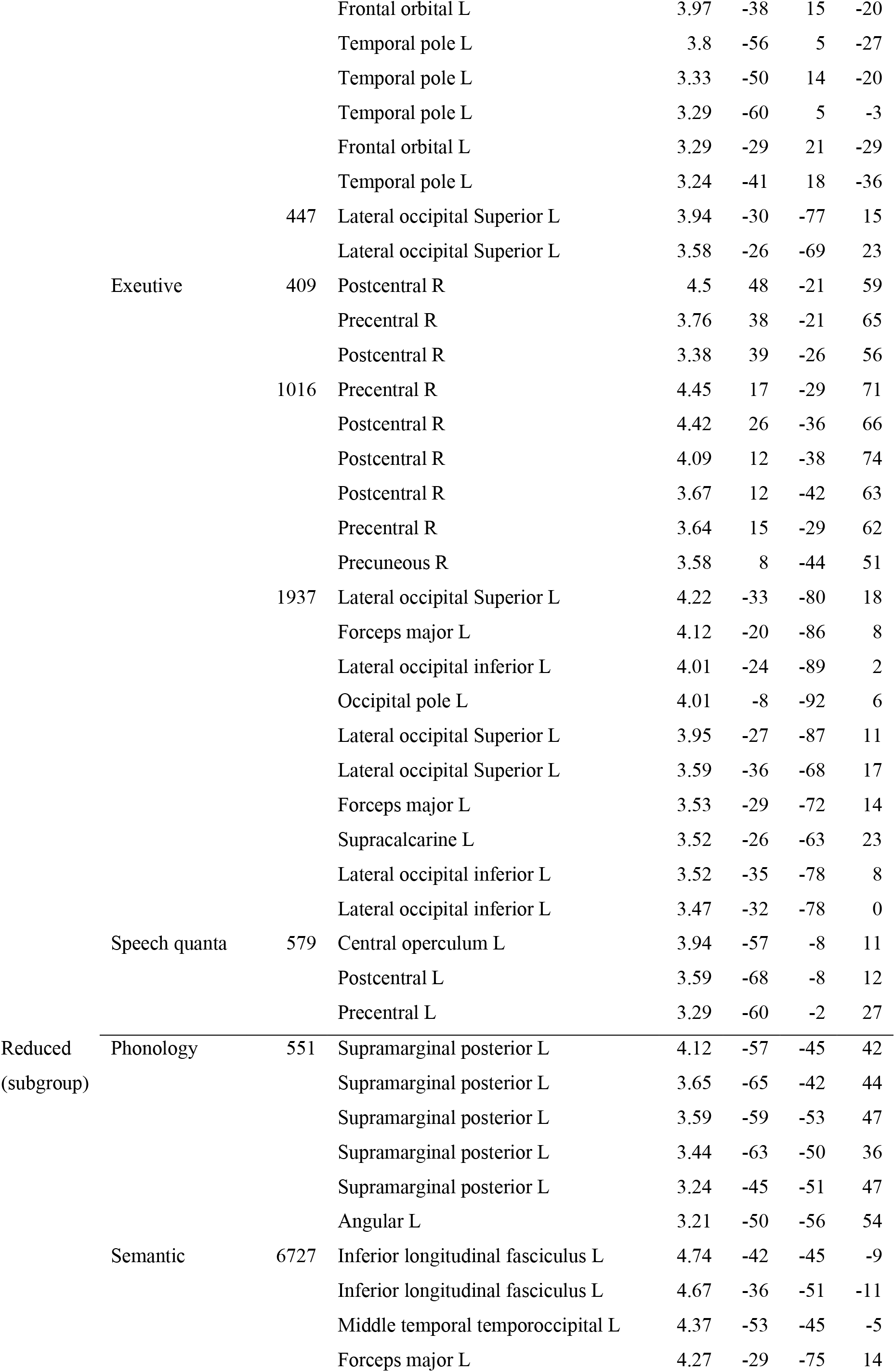
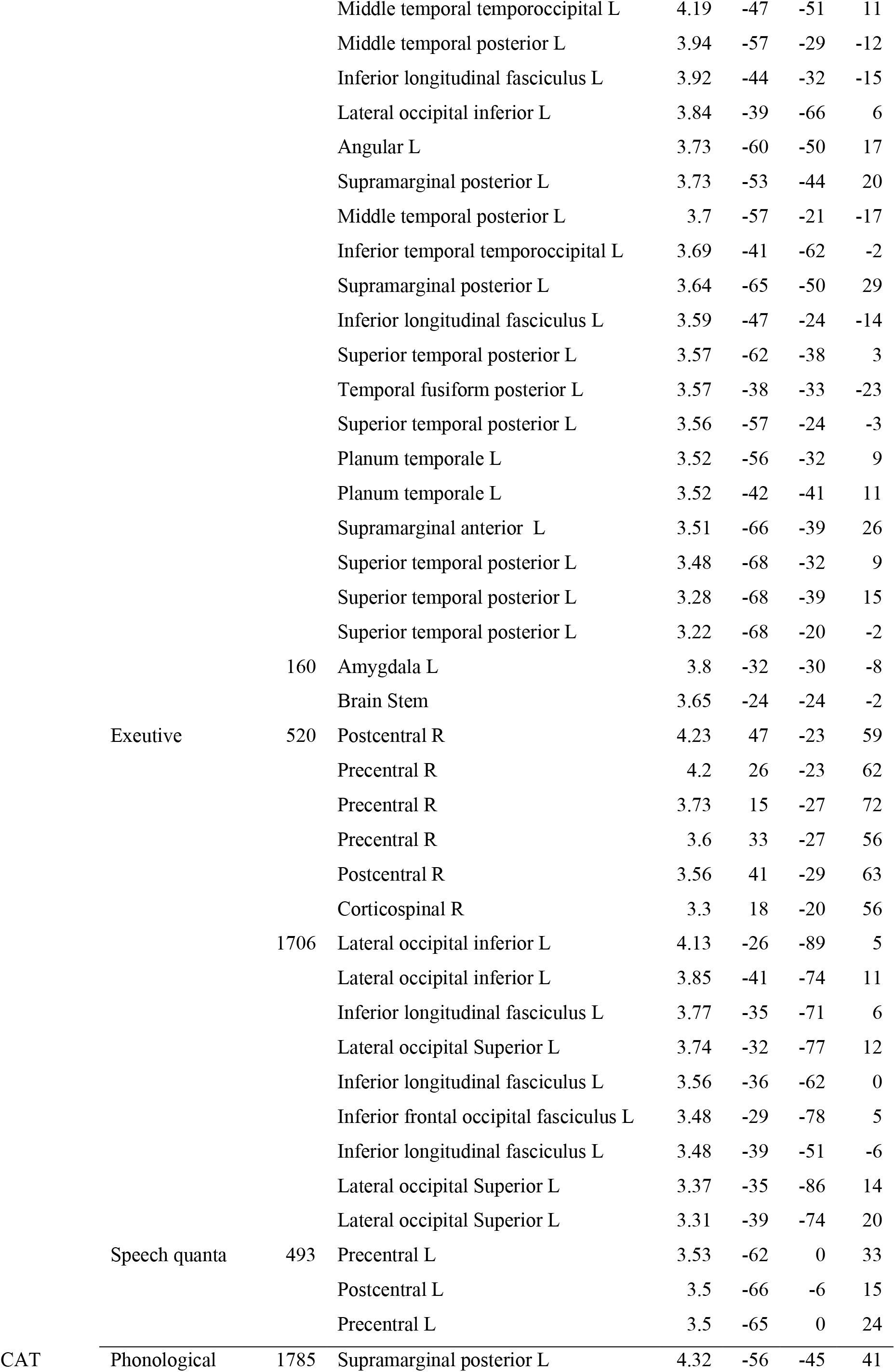
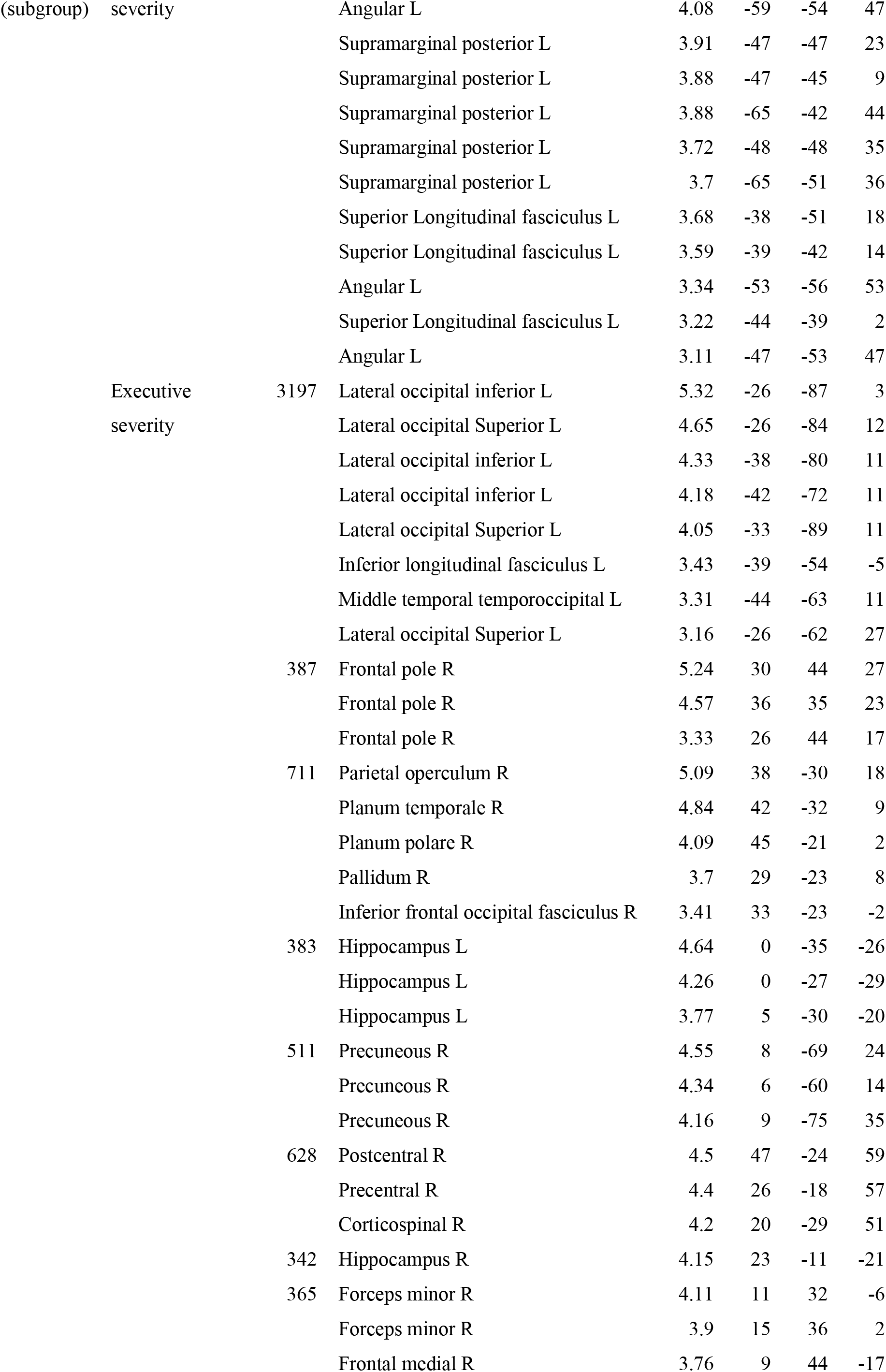
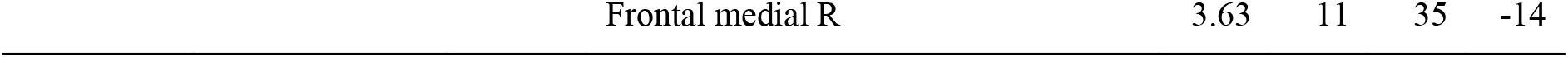
Neural correlates for PCA factors after accounting for lesion volume

### Section 5

**Figure 1.**
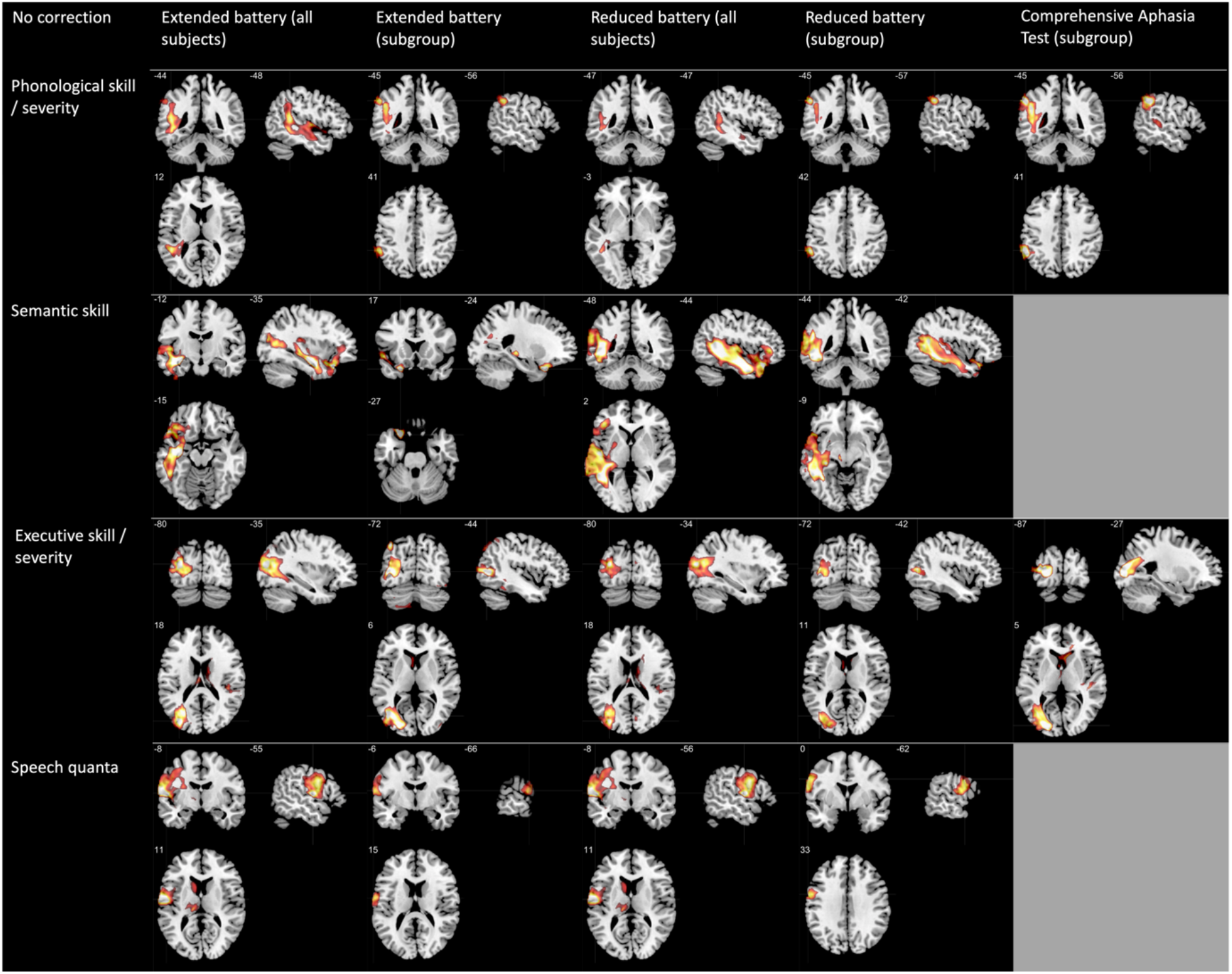
VBCM results for all principal components for each test battery. The components for each column were entered simultaneously and with no additional covariates. The results are thresholded using p < .001 voxelwise with family wise error cluster correction p < .05. The rows represent each principal component; phonological skill / severity, semantic skill, executive skill/severity and speech quanta. The grey patches in the final column indicate that there were no corresponding CAT components for semantic skill and speech quanta.

